# An Alpha-helical Lid Guides the Target DNA toward Catalysis in CRISPR-Cas12a

**DOI:** 10.1101/2022.09.05.506663

**Authors:** Aakash Saha, Mohd Ahsan, Pablo R. Arantes, Michael Schmitz, Christelle Chanez, Martin Jinek, Giulia Palermo

## Abstract

CRISPR-Cas12a is a powerful RNA-guided genome-editing system, also emerging as a robust diagnostic tool that cleaves double-stranded DNA using only the RuvC domain. This opens an overarching question on how the spatially distant DNA target strand (TS) traverses toward the RuvC catalytic core. Here, continuous tens of microsecond-long molecular dynamics and free-energy simulations reveal that an ⍺-helical lid, located within the RuvC domain, plays a pivotal role in the traversal of the TS by anchoring the crRNA:TS hybrid and elegantly guiding the TS toward the RuvC core, as also corroborated by DNA cleavage experiments. In this mechanism, the REC2 domain pushes the crRNA:TS hybrid toward the core of the enzyme, while the Nuc domain aids the bending and accommodation of the TS within the RuvC core by bending inward. Understanding of this cardinal process in the functioning of Cas12a will enrich fundamental knowledge and facilitate further engineering strategies for genome-editing.

---

Bacteria and other prokaryotes deploy an arsenal of CRISPR-Cas systems (<u>c</u>lustered <u>r</u>egularly <u>i</u>nterspaced <u>s</u>hort <u>p</u>alindromic <u>r</u>epeats, CRISPR-associated proteins) as a part of their adaptive immune response against foreign nucleic acid invasion.^1^ The CRISPR-associated proteins Cas9^2^ and Cas12a,^3^ being RNA-programmable endonucleases, launched unparalleled genome-editing applications across different fields of biotechnology, agriculture and medicine.^4,5^ The Cas12a nuclease came to the forefront with innovative applications such as extraordinarily rapid nucleic acid detection,^6^ also harnessed for the detection of the SARS-CoV-2.^7,8^ At the molecular level, Cas12a cleaves double-stranded target DNA using a single catalytic domain.^9–14^ Hence, conformational changes of the protein are critical for the accommodation and cleavage of DNA, but the mechanism of such a conformational transition is highly elusive.

Structures of Cas12a from *Francisella novicida* (FnCas12a) revealed a bilobed architecture, comprising of a recognition (REC) and a nuclease (NUC) lobe, flanked by the wedge (WED) domain (Fig. 1).^9–12^ Two alpha-helical domains (REC1 and REC2) constitute the REC lobe and mediate nucleic acid binding, while the NUC lobe comprises the RuvC catalytic domain and an adjacent Nuc domain. During its biophysical function, Cas12a uses CRISPR RNA (crRNA) for highly specific molecular recognition^15–17^ and cleavage of complementary DNA sequences. Upon recognition of a short protospacer-adjacent motif (PAM) by Cas12a, the crRNA binds one DNA strand (i.e., the target strand, TS) and forms a crRNA:TS hybrid, while the other non-target strand (NTS) gets displaced. Cas12a then confers double-stranded DNA breaks by using only the RuvC domain as its molecular scissor. This is strikingly different from Cas9, where two catalytic domains – HNH and RuvC – perform cleavage of the TS and NTS, respectively.^18^ This difference alludes to a conundrum on how a single nuclease domain could enable cleavage of both DNA strands in Cas12a. Indeed, while the NTS accommodates within the RuvC catalytic cleft, the TS stays spatially distant from the catalytic pocket, close to the REC lobe (Fig. 1). Experimental observations have inferred that the TS gets cleaved post the cleavage of the NTS.^9,12,19^ Together, this indicates a conformational change after the NTS cleavage, allowing the accommodation of the TS within the RuvC catalytic pocket. Nevertheless, the molecular details of these intriguing biophysical processes have remained incompletely understood.

**Fig. 1.**
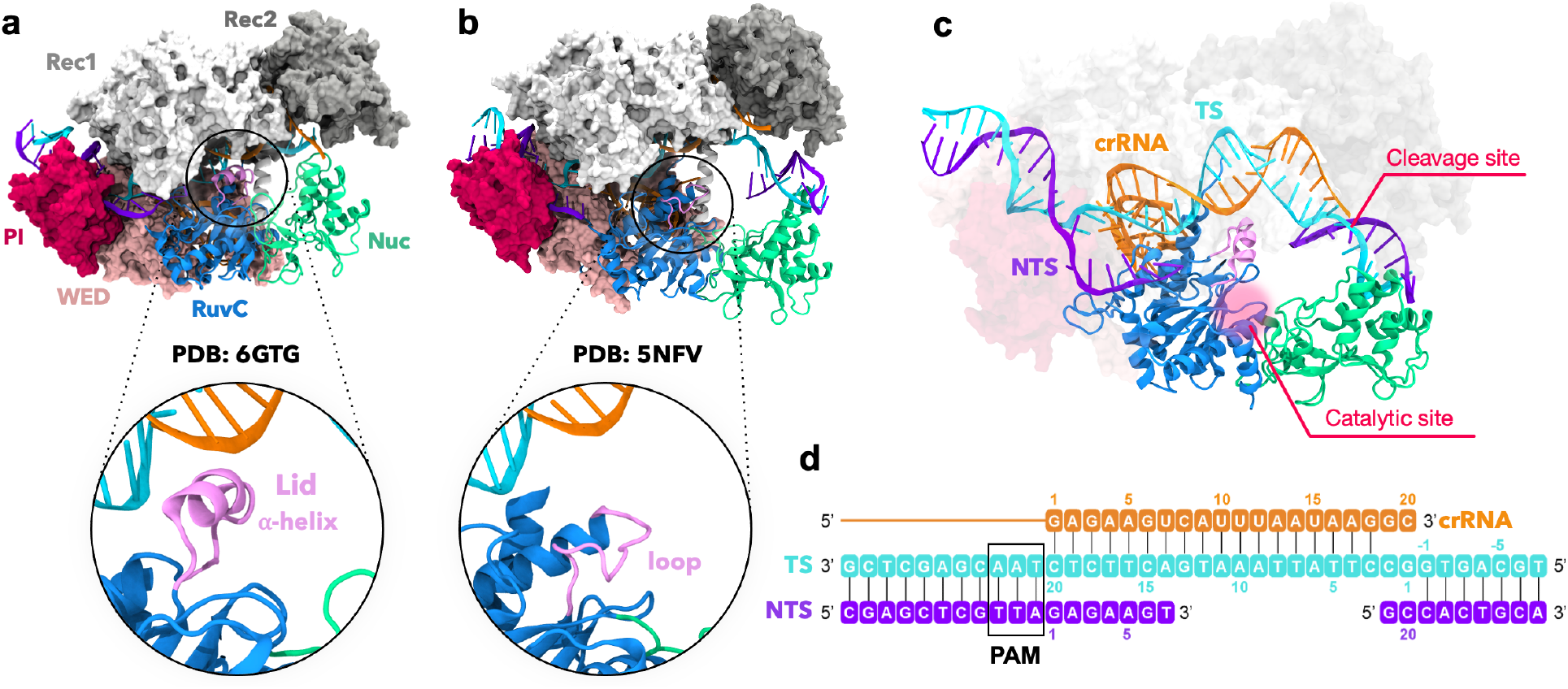
Structures of CRISPR-Cas12a after DNA non-target strand cleavage. The Cas12a protein is shown in molecular surface, highlighting the RuvC (blue) and Nuc (green) domains as ribbons. The crRNA (orange), the DNA target strand (TS, cyan) and non-target strand (NTS, violet) are also shown as ribbons. **a.** The cryo-EM structure (PDB: 6GTG)^12^ contains a short TS without the scissile phosphate. This structure displays an ⍺-helical “lid”, opening the RuvC domain (close-up view). **b.** The X-ray structure (PDB: 5NFV)^9^ reports a longer TS including the scissile phosphate and reconciling with the NTS. Here, the lid is unstructured (close-up view) and reconstructed through homology modelling. **c.** A more complete CRISPR-Cas12a modelled from the cryo-EM (a) and X-ray (b) structures, containing the ⍺-helical lid and a longer DNA TS including the cleavage site. **d.** Schematic representation of the nucleic acids.

Alongside biochemical and structural studies, single-molecule experiments^12,16,19–23^ have substantially contributed toward delineating the biophysical functioning of the Cas12a enzyme. The RuvC domain is structurally stable in Cas12a, while the REC2 and Nuc domains display increased flexibility and concerted motions upon DNA binding, which could favour the conformational transition of the TS toward the catalytic site. Such cooperative movements of REC2 and Nuc have been demonstrated by cryo-EM and single-molecule FRET experiments,^12^ and shown through molecular simulations.^24^ However, the protein dynamics and the conformational transitions orchestrating the traversal of the TS from the REC lobe to the RuvC catalytic site are yet to be identified. This is a critical question, understanding of which can lay the ground for further engineering toward improved genome editing.

Here, extensive Molecular Dynamics (MD) and free-energy simulations of Cas12a are used to unravel the process of DNA TS traversal and its repositioning within the RuvC catalytic core from a total simulation time of ~160 μs. Using Anton2, one of the fastest supercomputers available for MD simulations,^25^ we performed continuous tens of μs-long runs, identifying an ⍺-helical “lid” as a key structural element, and the REC2 and Nuc domains as the major players orchestrating the conformational transition. Next, free-energy methods captured the elaborate mechanism by which the ⍺-helical lid guides the DNA TS toward the catalytic core, as also supported by DNA cleavage experiments. Taken together, our findings shed light on a cardinal step in the functioning of Cas12a, which has previously remained highly elusive.

## Results

### An ⍺-helical lid anchors the crRNA:TS

Our investigations considered two structures obtained through cryo-EM and crystallographic techniques upon NTS cleavage with different lengths of DNA TS. The cryo-EM structure (PDB: 6GTG; EMD-0065,^12^ Fig. 1a) contains a short TS without the scissile phosphate. This structure has an open RuvC domain, displaying an ⍺-helical “lid” (residues L1008–K1021) that was held accountable for opening the RuvC domain for DNA accommodation.^12^ The X-ray structure (PDB: 5NFV,^9^ Fig. 1b) displays a full-length TS with the scissile phosphate, while the lid is structurally disordered.

Our ~10 μs long simulations of the 6GTG cryo-EM structure suggest that the ⍺-helical lid plays a crucial role in the major conformational changes of the system. Indeed, the ⍺-helical lid draws the TS toward itself by establishing extensive salt-bridge interactions between its Lys/Arg residues and the crRNA:TS hybrid backbone at PAM-distal sites (Supplementary Fig. 1). It is notable, however, that in this cryo-EM structure, the region of the TS downstream of the crRNA:TS hybrid is absent and consequently lacks the scissile phosphate.^12^ Hence, we modeled the downstream region of the DNA in the 6GTG structure based on the 5NFV X-ray structure,^9^ obtaining a more complete system of Cas12a after NTS cleavage (Fig. 1c). Two ~10 μs long replicates of this system consistently show that positively charged residues of the ⍺-helical lid interact with the crRNA:TS hybrid at PAM-distal sites after ~2 μs of MD (Fig. 2a-b). These include the stable interaction of K1018 with the TS at position 2, and R1014 with the crRNA at position 12. A bending of the TS at the end of the crRNA:TS hybrid is also observed that becomes more prominent after ~5 μs in both simulation replicates, leading to an ~40° bending with respect to the initial state (Fig. 2c). This is an important observation, which is in line with the experimental evidence of a distortion of the TS downstream of the crRNA:TS hybrid that was deemed necessary for it to reach the RuvC cleavage site.^26^ Remarkably, after ~5.8 μs of MD, the region of the TS including the scissile phosphate also moves toward the RuvC active site in both MD runs (Fig. 2d), further supporting the experimental indications. It is notable that biochemical and single-molecule experiments have shown that the cleavage site on the TS can mainly be located between base positions −2 and −4.^21,26^ Hence, we measured the distance between the scissile phosphate connecting the T_−2_ and G_−3_ nucleobases and the center of mass of the RuvC catalytic core (Fig. 2d, details in the Methods section). The RuvC catalytic core is composed of the D917, E1006, D1255 residues coordinating two Mg^2+^ ions, and displaying a conserved two-metal ion architecture similar to Cas9.^27,28^ During our ~10 μs long replicates, a substantial decrease of the computed distance indicates that the scissile phosphate approaches the RuvC core.

**Fig. 2.**
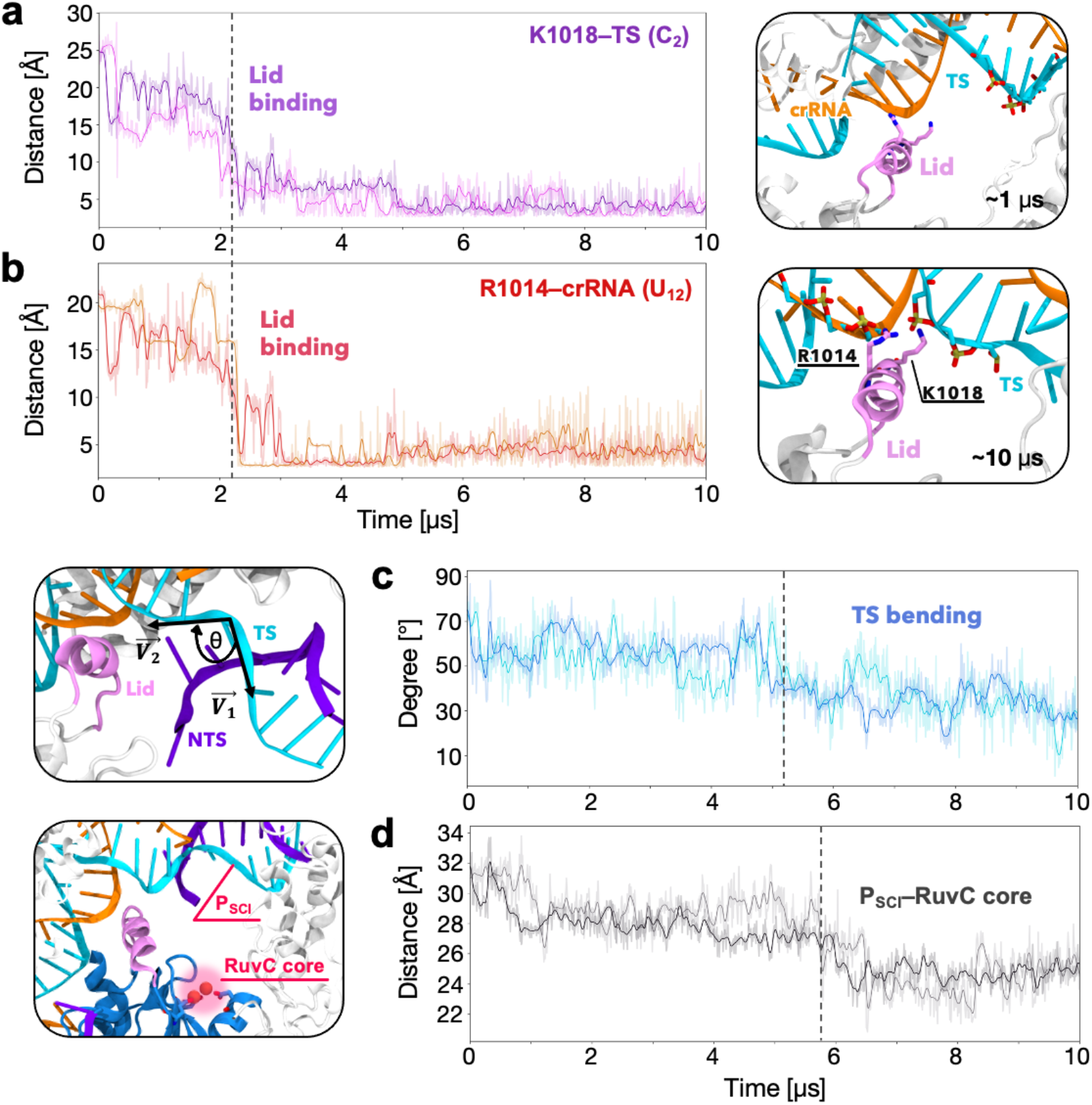
The ⍺-helical lid anchors the crRNA:TS hybrid. **a.** Interactions established by the positively charged residues of the ⍺-helical lid with the crRNA, and **b.** the DNA target strand (TS), evolving over ~10 μs of molecular dynamics simulations in two replicates. Representative snapshots on the right show the interactions between the ⍺-helical lid (mauve) and the crRNA:TS hybrid (crRNA: orange, TS: cyan) at different simulation times. **c.** Bending of the DNA TS, computed as angle *θ* between the vectors 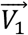 and 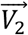 passing through the G_−1_, G_1_, C_2_, C_3_ and T_4_ (left panel, details in the Supplementary Methods). After ~5 μs of MD, the region of the TS at the end of the hybrid bends, leading to an ~40° bending with respect to the initial state. **d.** Distance between the scissile phosphate (P_SCI_, between the T_−2_ and G_−3_ nucleobases) and the center of mass of the RuvC catalytic core (left panel, see the Methods section). P_SCI_ moves toward the RuvC core after ~5.8 μs of MD. Dashed lines running through the graphs indicate critical conformational shifts. Data are reported for two simulation replicates of ~10 μs each.

Taken together, the two ~10 μs long simulations of Cas12a, including a longer TS and the scissile phosphate, confirm that the ⍺-helical lid anchors the crRNA:TS hybrid at PAM-distal sites. Upon these interactions, the TS bends and the scissile phosphate moves toward the RuvC core. The observed interactions thereby suggest that the lid could have a critical role in the traversal of the DNA TS for catalysis.

The 5NFV X-ray structure captured the protein complex after NTS cleavage, without a structured lid.^9^ Homology modeling reconstructed this disordered lid into an unstructured loop (Fig. 1b, see the Methods section). Molecular simulations (~10 μs in two replicates) of this system show a very high flexibility of the unstructured loop, especially in comparison with the ⍺-helical lid (Supplementary Fig. 2). Interestingly, this unstructured lid forms salt-bridges with the crRNA and intermittent interactions with the TS (Supplementary Fig. 3a-b). This, in turn, affects the bending of the TS and the movement of the scissile phosphate toward the RuvC catalytic site (Supplementary Fig. 3c-d). This indicates that the interactions of the lid with both the crRNA and TS are critical to achieve the bent conformation of the TS. Furthermore, this evidence bolsters the functional role of the ⍺-helical lid in the traversal of the TS toward the RuvC catalytic site.

### REC2 and Nuc facilitate the traversal of the TS

Our long-timescale simulations of CRISPR-Cas12a including a complete TS also reveal noteworthy conformational changes of the protein and the crRNA:TS hybrid, suggesting a cooperative dynamical mechanism fostering the traversal of the TS. To monitor these conformational changes, we analyzed the bending angles between the vector passing through the regions of interest and its coplanar perpendicular, with respect to the first frame of reference (details in the Supplementary Methods).

We observe that the long ⍺-helices of REC2 binding the crRNA:TS minor groove (residues 339-383, 474-543, 557-590; hereafter referred as REC2a) exhibit restricted motions. On the other hand, the short alpha helices connected by turns (residues 384-473, 543-557; referred to as REC2b) display a marked inward movement along the two ~10 μs long simulations. This is shown by a decrement in the bending angle of REC2b, which becomes more substantial after ~6 μs (Fig. 3a, gray and Supplementary Fig. 4a). This conformational change follows a similar trend of the arching of the crRNA:TS hybrid (Fig. 3a, orange). Indeed, the terminal major groove of the hybrid, adjacent to REC2b, arches toward the RuvC domain, as shown by the reduction of its vector angle in both simulation replicates (Supplementary Fig. 4b). The hybrid arching is remarkable ~6 μs of MD onward, alongside REC2b bending. Additionally, REC2b maintains extensive contacts with the crRNA:TS hybrid throughout the simulations (Supplementary Fig. 5). This indicates that the inward motion of REC2b helps the arching of the crRNA:TS hybrid. Both conformational changes are observed conspicuously upon the ⍺-helical lid binding to the crRNA:TS hybrid, occurring at ~2 μs in both simulation replicates (Fig. 2). These observations suggest the cooperativity between REC2 and the ⍺-helical lid. Indeed, the ⍺-helical lid anchors the crRNA:TS hybrid, which is followed by the bending of REC2, facilitating conformational changes of the hybrid toward the RuvC catalytic pocket.

**Fig. 3.**
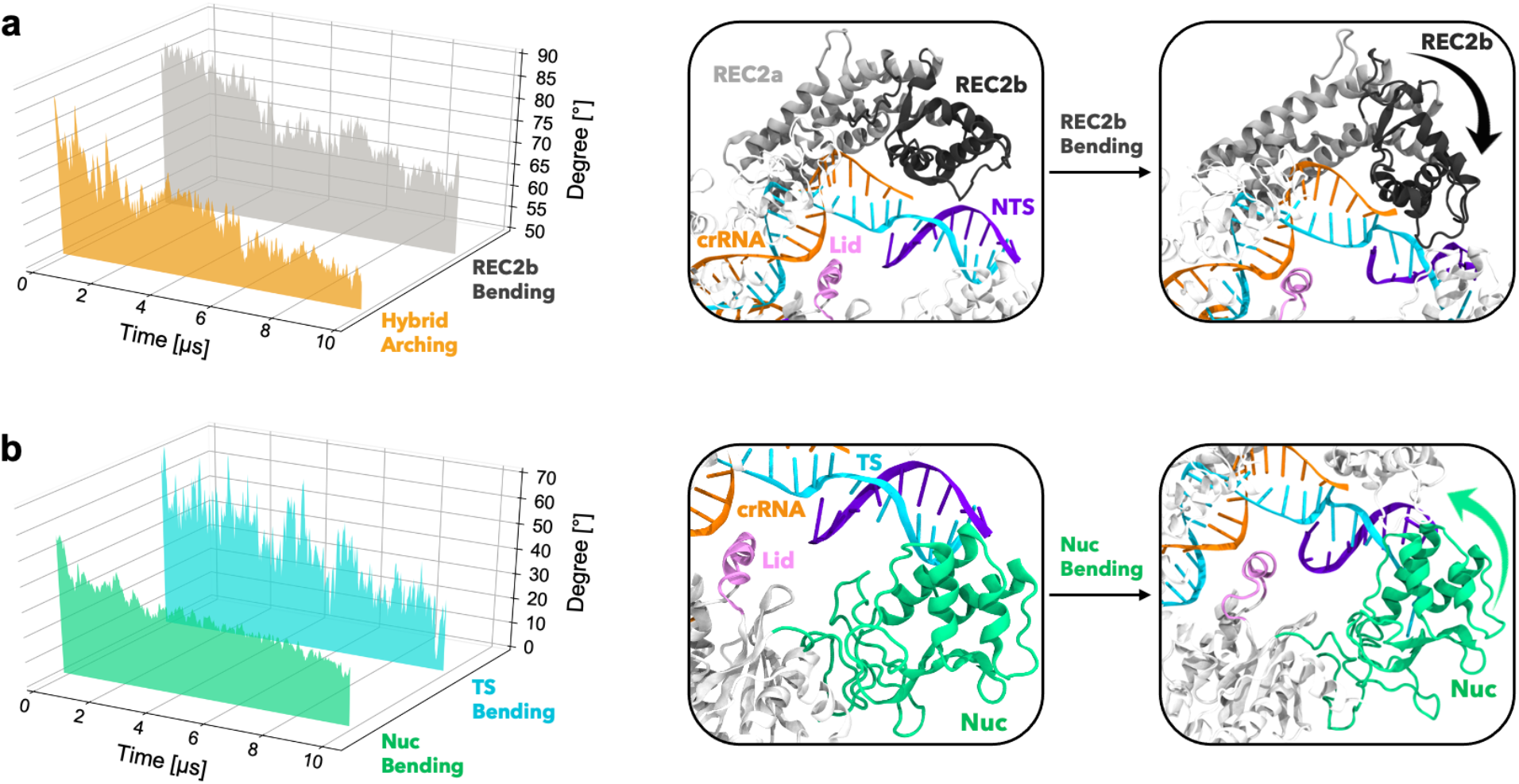
Bending of major domains in CRISPR-Cas12a. **a.** Bending of the REC2b region (residues 384-473, 543-557, gray) and arching of the terminal major groove of the crRNA:TS hybrid (base pairs G_19_:C_2_–U_15_:A_5_ orange) along ~10 μs of MD simulations of CRISPR-Cas12a including a complete TS. The REC2b and crRNA:TS hybrid conformational changes are shown on the right. **b.** Bending of the DNA TS (cyan, Fig. 2d) in concert with the bending of the Nuc domain (green) evolving over the multi-μs simulation. The panels on the right show the conformational changes. The curved arrows indicate the conformational changes in the domains. Details are in the SI Methods. Time evolution of the bending angles on 2-D plots is reported in Supplementary Fig. 4 for both simulation replicates.

Our simulations also show that the Nuc domain bends toward the core of the enzyme, as evidenced by the decrement of the Nuc bending angle (Fig. 3B, green). During this conformational transition, Nuc maintains a noteworthy number of contacts with the 5’-region of the TS (Supplementary Fig. 5). The DNA TS at the end of the hybrid also bends in both the simulation replicates (Fig. 3B, cyan and Supplementary Fig. 4). This similar trend of the Nuc and TS bending angles along with the persistent interactions between them suggest that the bending of Nuc aids in the bending of the DNA TS.

Cryo-EM and single-molecule experiments reported high flexibility and coupled dynamics of REC2 and Nuc, suggesting these conformational changes could be crucial for DNA cleavage.^12,16,20–23^ The continuous multi-μs long simulations reported here demonstrate that REC2 and Nuc bend toward the core of the enzyme (Fig. 3), while establishing multiple contacts between them (Supplementary Fig. 5). These largescale coupled motions exhibited by the major domains, along with the lid anchoring the TS, offer a compact conformation to the whole protein, facilitating the traversal of the TS toward the catalytic pocket.

### Traversal of the DNA TS and accommodation in the RuvC catalytic pocket

The region of the TS including the scissile phosphate is significantly far from the catalytic core (~30 Å, Fig. 1), which opens an overarching question regarding its traversal and accommodation within RuvC. To investigate this phenomenon, we performed free-energy simulations, sampling the dynamics of the complex from the complete Cas12a structure (Fig. 1c) to a modeled catalytically competent state (Supplementary Fig. 6). The latter was based on related X-ray structures holding the DNA TS within the RuvC active site (see Supplementary Methods). ~10 μs long MD simulations of this system revealed that no further major conformational change happens in the long timescale, especially in the dynamics of REC2 and Nuc (Supplementary Fig. 7a). This is indeed an important point, considering that the 6GTG and 5NFV structures underwent conformational changes over ~6-7 μs of continuous MD (Figs. 2–3). Importantly, the simulations also show that the RuvC active site with the DNA TS is remarkably stable (Supplementary Fig. 7b-d).

We performed Umbrella Sampling (US) simulations,^29^ studying the traversal of the DNA TS along two reaction coordinates (RCs). Structural,^9–14^ single-molecule^12,16,20,21^ and computational^24^ studies revealed that conformational changes in the Cas12a protein are needed to conduct the traversal of the TS. Hence, the difference in root-mean-square deviation (RMSD) of the C⍺ atoms between the initial and the final states appealed as an appropriate RC (RC1) to capture the slowest dynamics involved in the conformational transition. Additionally, the second RC (RC2) was selected as the distance between the center of mass of the RuvC catalytic core and that of the TS region including the scissile phosphate (i.e., the DNA nucleobases at positions −2 to −4, as indicated by biochemical and single-molecule data).^21,26^ The conformational transition was sampled over ~50 μs. Upon removing the biases incurred by the restraints, the free energy landscape (i.e., the Potential of Mean Force, PMF) was described along its minimum free energy pathway (see the Methods section, Supplementary Figs. 8-10).

We observe that the free energy profile flows freely (i.e., barrierless) toward the first minimum (Fig. 4a, blue line) as the traversal of the DNA TS is fostered by the interactions of the ⍺-helical lid with the crRNA:TS hybrid. Indeed, the ⍺-helical lid anchors the hybrid with its positively charged residues, while interacting with the nucleobases using N1009. A phenylalanine residue (F1012) is also observed to establish stacking interactions with the nucleobases of the TS (with T_−2_ and G_−3_). These critical interactions with the lid residues guide the descent of the TS. Then, a ~3 kcal/mol barrier is observed corresponding to the TS repositioning within the catalytic core, and the closure of the Nuc domain toward the core of the enzyme (Fig. 4A and Supplementary Fig. 11), constituting the slowest step in the process. These observations suggest that Nuc is critical in aiding the accommodation of the DNA TS within the RuvC catalytic core. Accordingly, the K1206-R1218 ⍺-helix/β-sheet region of Nuc maintains persistent interaction with the 5’-tail of the DNA TS (Supplementary Fig. 12). Hence, Nuc could hold the TS within the RuvC active core for catalysis, as also supported by recent single-molecule studies.^21^ Moreover, mutational studies on the Nuc β-sheet region reported compromised TS cleavage activity.^9,11^ Taken together, Nuc is critical in stabilizing the DNA TS at the level of the RuvC catalytic pocket, aiding its accommodation for cleavage, as demonstrated by our long timescale simulations.

**Fig. 4.**
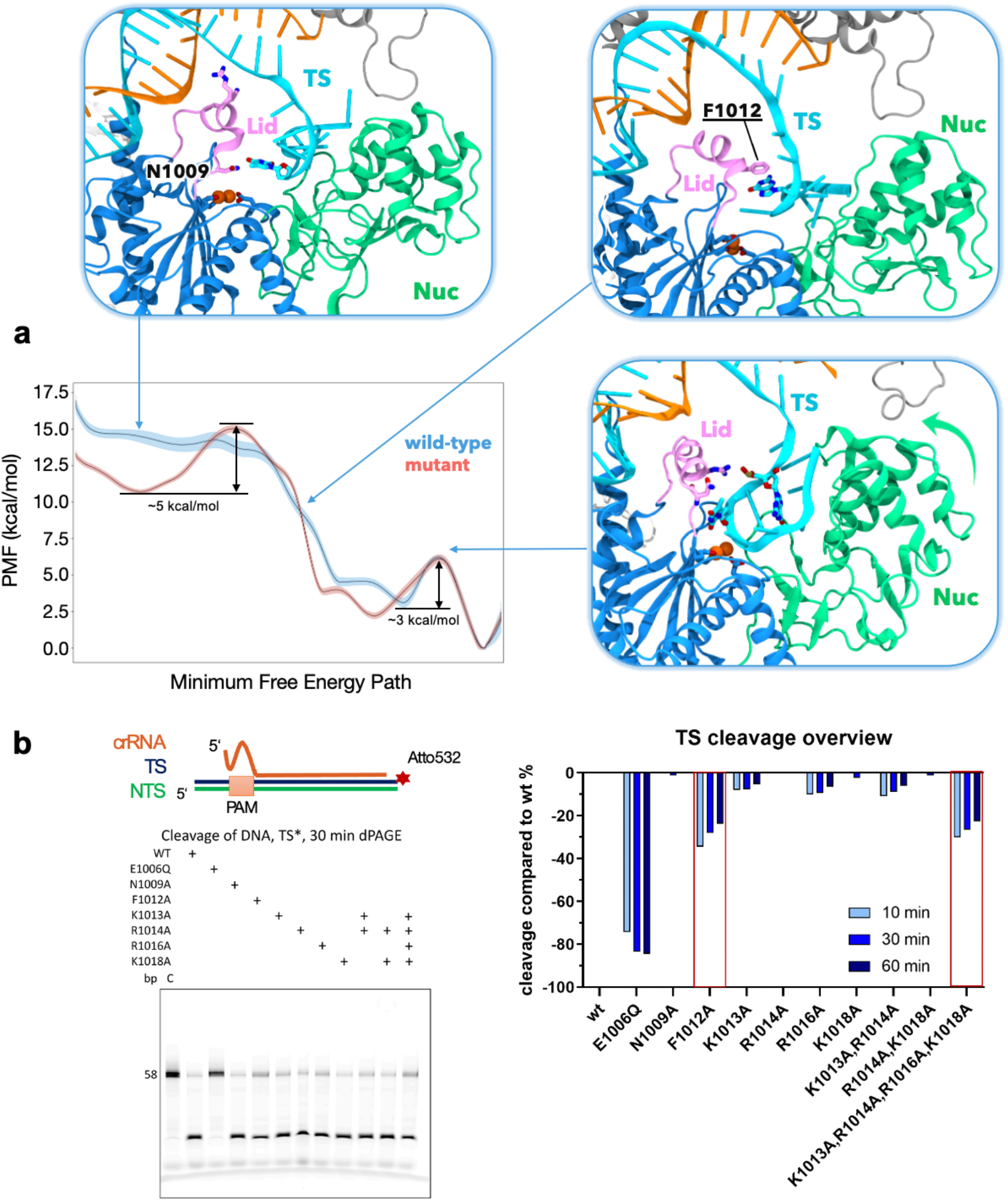
Traversal of the DNA target strand toward the RuvC catalytic core and role of the ⍺-helical lid. **a.** Free energy profiles for the traversal of the DNA target strand (TS) toward the RuvC core, in the wild-type WT Cas12a (blue) and upon mutating relevant residues of the ⍺-helical lid (F1012, N1009, K1013, R1014, R1016, K1018) into alanine (red). The Potential of Mean Force (PMF, in kcal/mol) is computed from 2-D umbrella sampling simulations and plotted along the minimum free energy path (see the Methods section). Representative snapshots along the free energy profile of the WT Cas12a are indicated by the arrows. Details for the Cas12a mutant are reported in Supplementary Fig. 10. **b.** In vitro DNA cleavage experiments, using Atto532 labelled 5’-end of TS, of Cas12a mutants showing a reduction in TS cleavage by the mutated lid systems compared to wild-type (WT) (left panel). Quantitation of cleavage activities show F1012A and the combined mutation (K1013A, R1014A, R1016A and K1018A) display noteworthy reduction in TS cleavage (right panel).

### Mutations of the lid adversely affect target strand traversal

To further verify the role of the lid in the traversal of the TS, we mutated the positively charged residues in the lid (K1013, R1014, R1016, K1018), along with F1012 and N1009, into alanine and we performed free energy simulations. The free energy profile was generated by collecting an ~50 μs ensemble, following the same US approach of the wild-type (WT) system.

The PMF of the mutated system shows an additional minimum with a free energy barrier of ~5 kcal/mol, which impedes a facile traversal of the TS (Fig. 4a, red line) as it was observed in the WT system.

This is because, in the mutated system, the TS locates more closely to the REC2 domain owing to the failure of interactions with the mutated lid (Supplementary Fig. 10). The PMF of the mutated system also shows a second free energy barrier of ~3 kcal/mol corresponding to the TS repositioning within the RuvC catalytic core and the closure of Nuc, similar to the WT system (Supplementary Fig. 11). This indicates that alanine mutations of the lid mainly affect the traversal of the TS, with a negligible impact on the repositioning of the TS within RuvC and the closure of Nuc.

To experimentally verify our observations, we introduced point mutations of residues in Cas12a that were identified through molecular simulations to facilitate TS traversal and assayed the TS cleavage activities of the mutant proteins in vitro (Fig. 4b). We observed substantial reduction in TS cleavage activities with combined alanine substitutions the positively charged residues (K1013A, R1014A, R1016A, K1018A) in the lid. In light of our simulations, this can be attributed to the loss of anchoring interactions between the lid and the crRNA:TS hybrid (Fig. 2). In contrast, single point mutations resulted in marginal reduction of the TS nuclease activity, most likely due to compensatory interactions with the hybrid by the adjacent positively charged residues in the lid. Notably, we observed a substantial reduction in TS cleavage with a single phenylalanine-to-alanine substitution (F1012A). The phenyl ring of F1012 was observed to stack with the nucleobases during our simulations of the TS traversal (Fig. 4A), suggesting that F1012 plays a critical role in guiding the TS toward the RuvC core. Taken together, our computational simulations and experimental validation establish a pivotal role of the ⍺-helical lid in the traversal of the TS toward the RuvC catalytic pocket.

## Discussion

In the CRISPR-Cas12a genome-editing system, the presence of a single RuvC catalytic site raises the question on how the enzyme could sequentially perform cleavages of the DNA NTS and TS respectively.^9,12,19^ Structures of Cas12a show that the TS stays spatially distant from the RuvC catalytic core, close to the REC lobe (Fig. 1). This implies that, following the NTS cleavage, a conformational change would allow the accommodation of the TS within the RuvC catalytic pocket, but the molecular details of this process have remained elusive.

Here, we performed extensive MD simulations of Cas12a after NTS cleavage, collecting an overall ensemble of ~160 μs. The simulations reveal a mechanism for the traversal of the DNA TS toward the RuvC catalytic core, in which an ⍺-helical lid plays a pivotal role. Structural studies showed that this ⍺-helical lid occludes the RuvC catalytic core in the binary RNA-bound Cas12a complex and opens for the accommodation of the NTS within the catalytic site.^12,30^ Multi-μs long MD simulations of Cas12a after NTS cleavage show that the ⍺-helical lid establishes extensive interactions with the crRNA:TS hybrid in its open conformation (Fig. 2a-b). These interactions involve the positively charged residues, including R1014 and K1018, that anchor the crRNA:TS hybrid at PAM-distal sites. A bending of the DNA TS at the end of the hybrid is also observed, followed by the TS region including the scissile phosphate moving toward the RuvC site (Fig. 2c-d). This observation is in agreement with experimental findings of a distortion of the DNA TS, downstream of the crRNA:TS hybrid, which was indicated to be necessary for it to reach the RuvC cleavage site.^26^

The REC2 domain also displays significant flexibility.^12,21,24^ Indeed, the short alpha helices connected by turns (referred here as REC2b) show prominent inward bending, which occurs in concert with the arching of the terminal major groove of the crRNA:TS hybrid (Fig. 3A). This suggests that REC2b, being heavier, pushes the PAM-distal major groove of the hybrid toward the inner core of the protein. Both conformational changes occur concertedly after the binding of the ⍺-helical lid to the hybrid (Figs. 2–3, and Supplementary Fig. 4). This indicates that the binding of the ⍺-helical lid prompts the arching of the hybrid, induced by REC2b pushing inward. Thus, the cooperativity between REC2 and the ⍺-helical lid could foster the traversal of the DNA TS toward the RuvC core. The Nuc domain also bends inward, in concert with the bending of the DNA TS (Fig. 3B). As REC2b and Nuc bend, they also significantly increase their interactions along the long timescale dynamics (Supplementary Fig. 5), resulting in a closed conformation of the complex. In this respect, cryo-EM and single-molecule experiments revealed high flexibility and coupled dynamics of REC2 and Nuc,^12^ also suggesting an open-to-close conformational change of the two domains.^20,22,23^ Single-molecule experiments further indicated such conformational changes to be crucial for DNA cleavage.^12,16,21^ Considering this evidence, the synergistic conformational changes of the major domains observed here, along with the anchoring of the ⍺-helical lid, poise the TS to traverse toward the RuvC active site for cleavage.

To elucidate the mechanism of DNA TS traversal toward the RuvC catalytic core, we performed free energy simulations. We found that the traversal of the TS occurs barrierless toward a first free energy minimum (Fig. 4a). This free-flowing profile is fostered by the interactions of the ⍺-helical lid with the hybrid. Indeed, positively charged residues of the lid establish extensive salt-bridge interactions with the TS, while F1012 stacks the nucleobases aiding the TS traversal. Importantly, mutations of the residues in the lid interacting with the hybrid impede a facile traversal of the TS with a ~5 kcal/mol barrier (Fig. 4a). Furthermore, our biochemical experiments also revealed that alanine mutations of these interacting residues adversely impact the cleavage of the DNA TS (Fig. 4b). While the combined mutations of the positively charged residues in the lid (K1013A, R1014A, R1016A, K1018A) result in a notable reduction of TS cleavage, single point mutation of F1012 alone reduces TS cleavage by ~40 %. Taken together, the lid interactions observed during our simulations establish a pivotal role of the ⍺-helical lid that guides the TS toward the RuvC core. In line with these observations, truncation of the lid has also shown to hamper target DNA cleavages.^12^ Moreover, a structural study on Casφ, a miniature Cas protein phylogenetically related to Cas12a, revealed a similar ⍺-helical lid exposing the RuvC domain for cleavage.^31^ Alanine substitution of the lid residues in this Casφ protein abolished double-stranded DNA cleavages, thereby underscoring the importance of the lid in the functioning of the enzyme.

Free-energy simulations also reveal that the Nuc domain is critical in aiding the accommodation of the TS within the RuvC core. Indeed, a ~3 kcal/mol barrier corresponds to the closure of the Nuc domain and further repositioning of the TS within the core of the enzyme (Fig. 4a). This region of Nuc maintains persistent interactions with the 5’-tail of the DNA TS and mutational studies of this region adversely impact TS cleavage.^9,11^ This suggests that Nuc could hold the TS at the RuvC site for cleavage, as also hinted by single-molecule experiments.^21^ Furthermore, free energy simulations of Cas12a including a mutated lid show that the closure of Nuc remains unaffected, and mutations in the lid rather hamper only the traversal of the DNA TS. Therefore, our findings indicate that while the lid is critical for the traversal of the DNA TS, Nuc can be decisive in the accommodation and repositioning of the TS in the RuvC catalytic core.

Taking together our findings and the available cryo-EM, biochemical and single-molecule experimental data, we propose a model for the traversal of the DNA TS strand toward the RuvC core for cleavage (Fig. 5). In this model, the ⍺-helical lid holds a critical role, along with the conformational changes in the major domains. Upon NTS cleavage, the ⍺-helical lid remains in its open conformation and anchors the crRNA:TS hybrid, establishing stable salt-bridge interactions (Fig 5a-b).

**Fig. 5.**
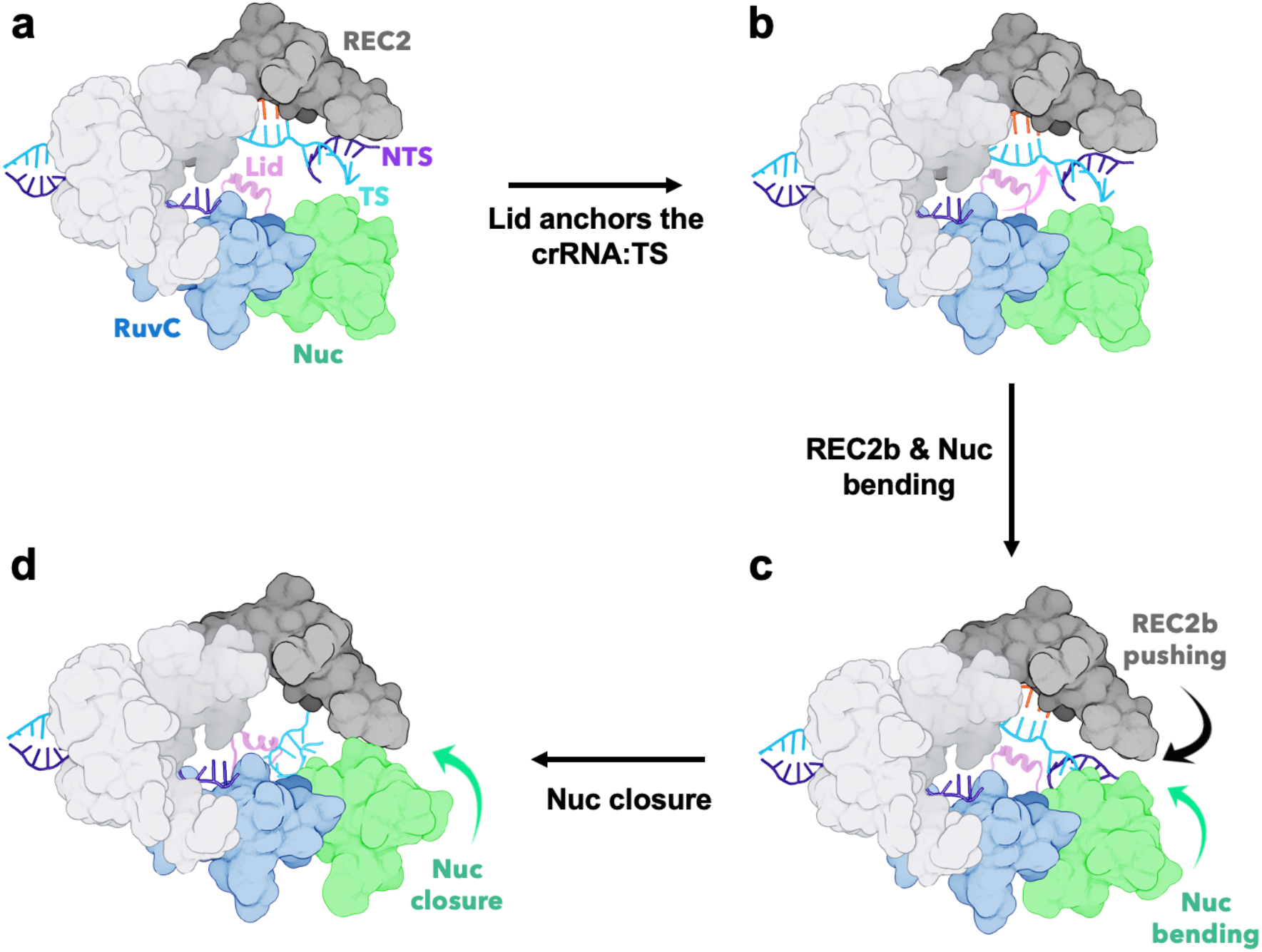
Model for the DNA target strand traversal toward the RuvC catalytic core in CRISPR-Cas12a. **a.** Upon cleavage of the DNA non-target strand (NTS), the α-helical lid assumes an open conformation, exposing the RuvC catalytic core, while the DNA target strand (TS) locates close to the REC2 domain.^12^ **b.** The α-helical lid, in its open conformation, anchors the crRNA:TS hybrid by establishing stable salt-bride interactions. **c.** The inward bending of the REC2 and Nuc domains pushes the crRNA:TS hybrid, while aiding the bending of the DNA TS and facilitating its conformational transition. **d.** The α-helical lid binds the DNA TS and guides it toward the RuvC catalytic core. The closure of Nuc aids the TS to accommodate within RuvC for cleavage.

Then, the REC2 domain pushes the terminal major groove of the hybrid through its REC2b region moving inward (Fig. 5c). As REC2b pushes the crRNA:TS hybrid, the Nuc domain simultaneously bends toward the core of the enzyme, aiding the bending of the DNA TS. These cooperative motions of the major domains prime the DNA TS for its traversal. The ⍺-helical lid now being in close proximity to the TS region around the scissile phosphate, establishes elaborate interactions with the TS and guides it toward the RuvC catalytic core. Finally, the closure of Nuc aids the TS to reposition and accommodate within the active site (Fig. 5d).

## Methods

### Structural Models

Molecular simulations were based on two structures of the *F. novicida* Cas12a obtained upon NTS cleavage: (1) the cryo-EM structure EMD-0065 (PDB: 6GTG)^12^ at 3.27 Å resolution and (2) the X-ray structure PDB: 5NFV,^9^ at 2.50 Å resolution. The 6GTG structure contains an ⍺-helical “lid” (residues L1008 – K1021, Fig. 1a), which is structurally disordered in the 5NFV structure. Hence, missing residues in the 5NFV structure were reconstructed through homology modeling using the SWISS-MODEL software (Fig. 1b).^32^ A third and more complete model system of Cas12a after NTS cleavage (Fig. 1c) was built starting from the 6GTG structure. Since this structure lacks the region of the TS downstream of the crRNA:TS hybrid, we modeled this region of the DNA based on the 5NFV X-ray structure. The catalytically competent state of Cas12a for TS cleavage was also considered, based on related crystal structures holding the DNA TS within the RuvC active site (see Supplementary Methods). All systems were embedded in explicit waters and counterions were added to neutralize the total charge, leading to periodic cells of ~138*149*167 Å^3^ and ~307,000 atoms for each system.

MD simulations were performed using the Amber ff19SB force field, with the ff99bsc0 corrections for DNA^33^ and the ff99bsc0+χOL3 corrections for RNA.^34,35^ The TIP3P model was employed for water,^36^ and the Li & Merz model was used for Mg^2+^ ions.^37^ We have extensively employed these force field models in computational studies of CRISPR-Cas systems,^38^ showing that they perform well for long timescale simulations on Anton-2.^39^ The Li & Merz model also reported a good description of Mg^2+^ bound sites, in agreement with quantum/classical simulations.^28,40^ An integration time step of 2 fs was employed. All bond lengths involving hydrogen atoms were constrained using the SHAKE algorithm. Temperature control (300 K) was performed via Langevin dynamics,^41^ with a collision frequency γ = 1. Pressure control was accomplished by coupling the system to a Berendsen barostat^42^ at a reference pressure of 1 atm and with a relaxation time of 2 ps. The systems were subjected to energy minimization, thermalization and equilibration as detailed in the Supplementary Methods. Then, ~120 ns of conventional MD were carried out in an NVT ensemble using the GPU-empowered version of AMBER 20.^43^ The well-equilibrated systems were used as starting points for simulations on Anton-2,^25^ a special-purpose supercomputer for multi-μs long MD simulations.

Long timescale MD simulations on Anton-2 were performed using the same force field parameters used for conventional MD simulations. A reversible multiple time step algorithm^44^ was employed to integrate the equations of motion with a time step of 2 fs for short-range nonbonded and bonded forces and 6 fs for the long-range nonbonded forces. Simulations were performed at constant temperature (300 K) and pressure (1 atm) using the multigrator integrator implemented in Anton-2.^45^ The k-Gaussian split Ewald method^46^ was used for long-range electrostatic interactions. Hydrogen atoms were added assuming standard bond lengths and were constrained to their equilibrium position with the SHAKE algorithm.^47^ By using this approach, each system was simulated on Anton-2, obtaining continuous MD runs of ~10 μs and in replicates. We obtained multiple trajectories of the 6GTG cryo-EM structure (simulated for ~10 μs) and its more complete model including a longer DNA (~10 μs in two replicates), as well as of the 5NFV X-ray structure (~10 μs in two replicates). The structural model of the catalytically competent Cas12a including the DNA TS within the RuvC site was also simulated for ~10 μs. This resulted in a total of ~60 μs of all-atom MD simulations.

### Umbrella Sampling simulations

The umbrella sampling (US) method^29^ was used to compute the free energy profile associated with the traversal of the DNA TS toward the RuvC active site. In this method, a number of simulations (US windows) are run in parallel with additional harmonic bias potential applied to selected Reaction Coordinates (*RCs*):

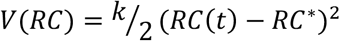

where *V*(*RC*) is the value of the bias potential, *k* is a bias force constant, *RC*(*t*) is the value of *RC* at given time *t* and *RC*\* is the reference value of *RC*. By using different *RC*\* values in each US window, one can sample the biased probability distribution *p_b_*(*RC*) along the whole *RC* range of interest. We performed two-dimensional (2-D) US simulations using two *RCs*. *RC*1 was the difference in root-mean-square deviation (RMSD) of the C⍺ atoms between the initial (Fig. 1c) and the final (Fig. S6d) states. *RC*2 was the distance between the center of mass of the RuvC catalytic core (D917, E1006, D1255) and the TS region including the scissile phosphate (i.e., the DNA nucleobases at positions −2 to −4, as indicated by biochemical and single-molecule experiments).^21,26^ *RC*1 was divided into bin sizes of 0.2 Å *RMSD* difference and each window was run for ~10 ns with a collective force constant *k* of 50 kcal mol^−1^ Å^−2^. *RC*2 was discretized into bin sizes of 0.2 Å and individual windows were run for ~15 ns with a collective force constant *k* was 50 kcal mol^−1^ Å^−2^. Overall, we simulated 28 (*RC*1) * 119 (*RC*2) = 3,332 windows, resulting in a collective ensemble of ~50 μs simulations.

Two independent sets of 2-D US simulations were performed (each for ~50 μs): (i) for the wild-type system and (ii) upon mutating relevant residues of the ⍺-helical lid (N1009, F1012, K1013, R1014, R1016, K1018) into alanine. Then, the free energy profiles were computed using the Weighted Histogram Analysis (WHAM) method^48^ upon removing the initial 1/3^rd^ of the US trajectories from each window, corresponding to the relaxation of the system. Analysis of the conformational ensembles was performed on the reweighted trajectories (see Supplementary Methods). Analysis of the free energy landscape was performed by computing the minimum free energy pathway using an approach similar to the Dijkstra algorithm.^49^ In this approach, the map is sampled from the origin point, propagating to the lowest energy point available on each iteration until it reaches the target point. This enabled us to consider both the *RCs* along a minimum free energy path. The error estimation on the minimum free energy pathways was performed using the Monte Carlo bootstrap error analysis.^50^ Full details on the 2-D US simulations are reported in the Supplementary Methods, including the convergence of the free energy profiles (Supplementary Fig. 8) and the 2-D free energy surfaces (Supplementary Fig. 9).

### FnCas12a expression and purification

The DNA sequence of *Francisella tularensis subsp. novicida* U112 (Fn)Cas12a (WP_003040289) was codon optimized for heterologous expression in Escherichia coli (E. coli) and synthesized by GeneArt (Thermo Fisher Scientific). The FnCas12a gene was inserted into the 1B plasmid (Addgene #29653) using ligation-independent cloning (LIC), resulting in a construct carrying an N-terminal hexahistidine tag followed by a tobacco etch virus (TEV) protease cleavage site. Point mutations were introduced by Gibson assembly of the PCR-amplified vector backbone with gBlock Gene Fragments (IDT) encoding the individual mutations. The sequences of the synthetic gene, primer and gBlocks are listed in Tab. 1. Mutant FnCas12a constructs were purified as for wild type. Purification of Cas12a was done as described previously.^51,52^ In brief, Cas12a constructs were expressed in E. coli BL21 Rosetta2 (DE3) cells (Novagen, Wisconsin, USA). Cells were lysed in 20 mM Tris pH 8.0, 500 mM NaCl, 5 mM imidazole, 1 μg/mL pepstatin, 200 μg/mL 4-(2-Aminoethyl)benzenesulfonyl fluoride hydrochloride (AEBSF) by ultrasonication. Clarified lysate was applied to a 10 ml Ni-NTA (Sigma-Aldrich) affinity column. The column was washed with 20 mM Tris pH 8.0, 500 mM NaCl, 5 mM imidazole, and bound protein was eluted by increasing imidazole concentration to 250 mM. Eluted protein was dialysed against 20 mM HEPES pH 7.5, 250 mM KCl, 1 mM DTT, 1 mM EDTA overnight at 4 °C in the presence of TEV protease to remove the 6xHis- affinity tag. Cleaved protein was further purified using a HiTrap HP Heparin column (GE Healthcare, Illinois, USA), eluting with a linear gradient to 1.0 M KCl. Elution fractions were pooled, concentrated, and further purified by size exclusion chromatography using a Superdex 200 (16/600) column (GE Healthcare) in 20 mM HEPES-KOH pH 7.5, 500 mM KCl, 1 mM DTT yielding pure, monodisperse proteins. Aliquots were flash-frozen in liquid nitrogen and stored at −80°C.

### FnCas12a nuclease activity assays

In vitro nuclease activity assays were conducted using purified WT or mutant FnCas12a proteins programmed with a crRNA targeting the λ- sequence (oMS017) and dsDNA substrates containing either fluorescently labelled TS (oDS285:oDS271) or NTS (oDS270:oDS288). FnCas12a and crRNA were mixed in a ratio of 1:1.2 and incubated for 10 min at 37 °C to allow binary complex formation. Reactions were started by addition of target dsDNA (FnCas12a:dsDNA, 10:1) and incubated at 37 °C. All samples were assembled in a final reaction volume of 20 μL containing 0.5 μM (mutant) FnCas12a, 0.6 μM crRNA and 50 nM dsDNA in a final buffer of 10 mM HEPES-KOH pH 7.5, 250 mM KCl, 5 mM MgCl2, 0.5 mM DTT. Reactions were stopped at indicated time points by addition of EDTA and Proteinase K (Thermo Fisher Scientific) in final concentrations of 80 mM and 0.8 mg/mL, respectively and incubated for 30 min at 37 °C. Samples were mixed with equal volume of a 2X dPAGE loading dye (95 % formamide, 25 mM EDTA, 0.15 % OrangeG), heated to 95 °C for 5 min and resolved on a 15 % denaturing (7 M Urea) polyacrylamide gels run in 0.5 X TBE buffer. Fluorescence of the ATTO532-labeled cleavage products was detected using a Typhoon FLA 9500 gel imager; the cleavage rate was quantified based on loss of uncleaved substrate DNA using ImageQuant TL v.8.2.0. Sequences of RNA and DNA oligos utilized in the nuclease assays are listed in Tab. 2.

## Acknowledgments

We thank Chinmai Pindi for useful discussions. This material is based upon work supported by the National Institute of Health (Grant No. R01GM141329, to GP) and the National Science Foundation (Grant No. CHE-2144823, to GP). MJ acknowledges support from the Swiss National Science Foundation (31003A_182567). Computer time for MD has been awarded by XSEDE under Grant No TG-MCB160059 and by NERSC under Grant No M3807 (to GP).

## Author Contribution

AS, MA and PRA performed molecular simulations and analysed data. ^†^MA and PRA contributed equally. MS and CC performed DNA cleavage experiments, supervised by MJ. GP conceived this research. AS and GP wrote the manuscript with critical input from all authors.

## Data Availability Statement

The datasets generated during and/or analysed during the current study are available from the corresponding author on reasonable request.

